# Uniquely high spontaneous mutational load in blood cells of XP-C patients

**DOI:** 10.1101/2025.07.16.665113

**Authors:** Gordon Fan-Huang, Elizabeth L. Schmidt, Moonsook Lee, Shixiang Sun, Christina Boull, Sheilagh M. Maguiness, Jan Vijg, Laura J. Niedernhofer, Alexander Y. Maslov

## Abstract

We discovered a uniquely high spontaneous somatic mutational load in peripheral blood mononuclear cells (PBMCs) of Xeroderma Pigmentosum group C (XP-C) patients, characterized by elevated single nucleotide variants associated with mutation signatures SBS8 and SBS32, as well as an enrichment of single-nucleotide cytosine deletions. This hypermutability was markedly lower in fibroblasts, suggesting a replication-dependent mechanism of mutagenesis due to deficient global genome nucleotide excision repair (GG-NER). Our findings reveal distinct molecular subtypes within XP defined by spontaneous mutational load in normal blood cells and demonstrate that the exceptionally high mutation burden in XP-C leukemia originates from mutations already present prior to malignant transformation.

## Introduction

Xeroderma Pigmentosum (XP) is a rare autosomal recessive disorder characterized by sensitivity to ultraviolet (UV) light, caused by germline mutations in seven DNA repair genes (XPA to XPG) or the bypass polymerase POLH/XPV. These mutations affect nucleotide excision repair (NER), which consists of two sub-pathways: global genome NER (GG-NER) and transcription-coupled NER (TC-NER)^1^. In GG-NER, XPC and XPE scan the entire genome for helical distortion, then recruit the repair machinery required for both sub-pathways, including XPA, XPB, XPD, XPF, and XPG. Consequently, mutations in XPC and XPE specifically impair GG-NER, while mutations in the other genes disrupt both GG-NER and TC-NER. This inability to remove UV-induced DNA adducts causes a ∼10,000-fold increased incidence of skin cancer in XP patients.^2^ XP complementation group C (XP-C) is distinct, as XP-C patients also have a ∼1,000-fold increased risk of hematological cancers.^3^

To investigate the molecular basis for these divergent clinical outcomes, we measured somatic mutation frequency in normal, non-tumor peripheral blood mononuclear cells (PBMCs) and primary fibroblasts from patients across different XP complementation groups and their unaffected first-degree relatives. Because somatic mutations are rare and occur stochastically across individual cells, they cannot be accurately quantified using conventional bulk sequencing.^4^ To overcome this limitation, we applied our recently developed error-corrected next-generation sequencing assay (ecNGS) based on single-molecule sequencing (SMM-seq), enabling highly sensitive and quantitative detection of low-abundance mutations.^5^

Using this approach, we reveal that GG-NER deficiency in XP-C blood cells leads to pronounced, replication-associated accumulation of somatic mutations. Notably, the mutational patterns observed in normal hematopoietic cells closely mirror those seen in XP-C leukemias, providing direct molecular evidence that a cancer-like mutational burden can arise prior to malignant transformation. Our results establish a mechanistic link between GG-NER deficiency, replication-dependent mutagenesis, and cancer predisposition, and offer a new conceptual framework for understanding how somatic mutation accumulation in inherited genome instability syndromes drives cancer risk.

## Results and Discussion

The analysis revealed an approximately 10-fold higher burden of somatic single nucleotide variants (SNVs) in blood from XP-C patients as compared to blood from XP-C heterozygous controls (Figure 1A). Strikingly, the SNV load did not significantly differ between XP-A, XP-D, or XP-F patients and their respective controls. The increased SNV frequency in XP-C blood samples predominantly reflected mutation signatures SBS8, known to be associated with replication,^6^ and SBS32, a signature not previously linked to XP (Figure 1B). Additionally, XP-C patient blood exhibited a significantly elevated frequency of small INDELs (Figure 1C) which were almost exclusively 1-nucleotide deletions of cytosine (Supplementary Figure 1). SMM-seq analysis of fibroblasts from the same XP-C subjects also revealed a statistically significant but modest (2.2-fold) increase in somatic mutations. However, no significant changes in INDEL frequency were observed in fibroblasts from XP-C or any other XP complementation groups (Supplementary Figure 2).

**Figure 1.**
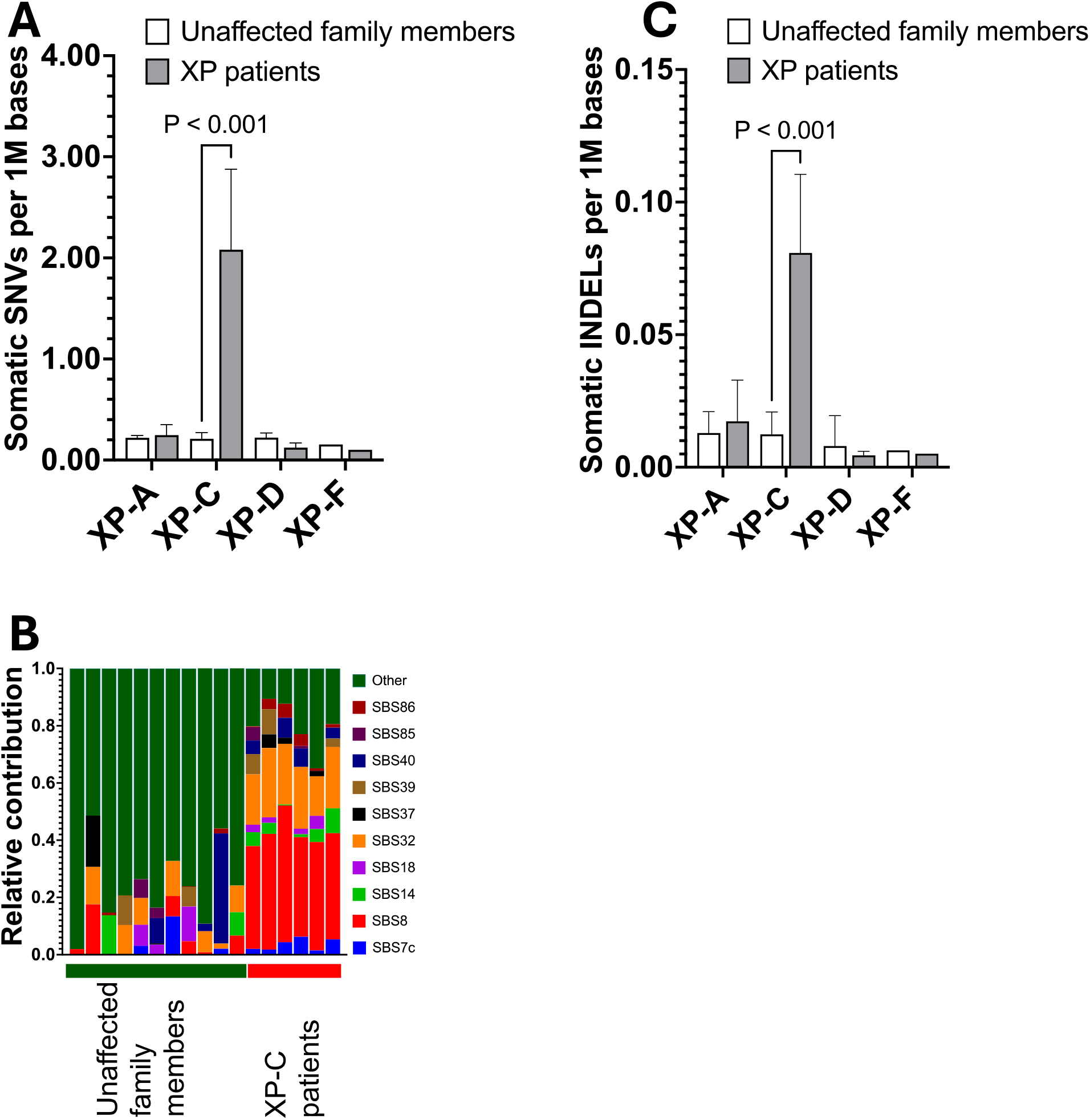
Somatic Mutation Burden and Spectra in Xeroderma Pigmentosum (XP) Patients Across Different Complementation Groups. Peripheral blood mononuclear cells (PBMCs) from XP-C patients exhibit significantly higher spontaneous somatic SNV (**A**) and INDEL (**C**) loads compared to healthy controls and other XP complementation groups. SNV frequencies were compared between XP patients from different complementation groups and unaffected first-degree relative controls. Data are presented as mean ± SD. (**B**) – Somatic mutation signatures identified in peripheral blood from XP-C patients and heterozygous carriers.

Our findings suggest two molecular subtypes of XP: one with low spontaneous SNV loads in non-sun-exposed somatic cells (e.g., XP-A, XP-D, XP-F), which is prone to developing UV-induced skin tumors, and another with high spontaneous SNV frequency in non-sun-exposed somatic cells (e.g., XP-C), which, in addition to skin cancer, is at higher risk of developing internal tumors, primarily hematological.^3^

A plausible explanation for the low spontaneous mutational load in XP complementation groups with defects in both GG-NER and TC-NER is that when TC-NER is absent, persistently stalled transcription complexes induce apoptosis, averting the generation of mutant cells.^7^ This may explain the progressive neurological degeneration and extreme photosensitivity observed in these groups.^8^ In contrast, when only GG-NER is absent, transcription blockage at unrepaired lesions is less likely to occur because TC-NER can still repair the damage by detecting lesions via stalled RNA polymerase II during transcription and recruiting the intact repair machinery to remove the blockage. As a result, cells survive despite the absence of GG-NER. However, unrepaired DNA adducts in non-transcribed regions persist and can become fixed into mutations during replication by error-prone translesion polymerases that also frequently introduce single-base slippage errors.^9^ This results in a highly excessive mutational load in GG-NER-deficient cells, as seen in XP-C patients and model systems.^10^ The particularly high mutation burden observed in XP-C PBMCs compared to syngeneic fibroblasts is therefore likely attributable to their much higher replication rate and turnover, amplifying the effects of this replication-dependent mutagenic process.^11^

Notably, leukemic cells from XP-C patients were reported to have a 25-fold increase in somatic mutations compared to sporadic cases of leukemia,^3^ aligning with our data. Moreover, the mutation spectra in normal blood cells of XP-C patients (Supplementary Figure 3A), characterized by a higher prevalence of C-to-A and A-to-T mutations, closely resembled those in XP-C leukemic cells (cosine similarity: 0.981; Supplemental Figure 3B), which also exhibited signature SBS8 as well as frequent 1-nucleotide deletions of cytosine.^3^ These striking similarities indicate that the extreme mutational burden characteristic of XP-C leukemia likely originates from mutations already present in normal cells before malignant transformation.

Our study is the first to quantify somatic mutation rates in normal human tissues across XP complementation groups, using XP as a model for inherited DNA repair deficiency. Critically, we show that GG-NER loss in XP-C blood cells drives replication-dependent accumulation of somatic mutations, establishing a pre-cancerous mutational burden that likely underlies the high incidence of internal, particularly hematologic, cancers in XP-C. These insights could inform new therapeutic strategies aimed at modulating cell death and DNA repair pathways, with the potential to benefit patients with XP and other DNA repair deficiency syndromes at increased risk for cancer.

## Methods

### Patient Recruitment and Sample Collection

Patients diagnosed with Xeroderma Pigmentosum (XP) and their unaffected family members were recruited through the University of Minnesota Medical School. Inclusion criteria for XP patients included a confirmed clinical diagnosis, while unaffected relatives were identified based on the absence of clinical symptoms and, where available, genetic confirmation. All participants provided written informed consent prior to enrollment, in accordance with ethical guidelines approved by the University of Minnesota Institutional Review Board. Blood samples were collected using standard venipuncture techniques. Samples were processed immediately to separate plasma, serum, and peripheral blood mononuclear cells (PBMCs), which were stored at –80°C until analysis. Recruitment strategies included direct physician referrals and outreach through patient advocacy groups to ensure adequate representation of the study population.

### DNA Extraction, Library Preparation, and Sequencin

DNA was extracted from frozen PBMCs using the Quick-DNA Miniprep Kit (Zymo Research), quantified with a Qubit fluorometer (Thermo Fisher), and assessed for quality using the TapeStation system (Agilent Technologies). Sequencing libraries for genotyping were prepared using the NEBNext Ultra II DNA Library Prep Kit for Illumina (New England Biolabs) following the manufacturer’s recommendations. SMM-seq libraries were prepared as previously described.^5^ Sequencing was performed by Novogene (USA) on the NovaSeq X Plus platform in 150 bp paired-end mode.

### Variant Calling and Data Analysis

Sequencing reads were aligned to the human reference genome (hg19), and variant calling was performed as previously described.^5^ VCF files were processed using the MutationalPatterns software package (v3.16.0).^12^ To characterize underlying SNV mutational signatures, the optimal linear combination of COSMIC v3.2 signatures was determined for each sample individually using non-negative least squares fitting. To compare signatures observed in normal XP-C PBMCs with those previously reported in XP-C leukemia, publicly available data from Yurchenko et al. (2020)^3^ were utilized. Mutations from normal XP-C PBMCs and XP-C leukemia samples were pooled separately to generate two group-level mutational profiles. Signature composition was calculated for each group, and cosine similarity was used to assess how closely the XP-C PBMC signature resembled the XP-C leukemia signature. GraphPad Prism version 10 was used for all statistical analyses; statistical significance was determined using a two-tailed t-test.

### Declaration of generative AI and AI-assisted technologies in the writing process

During the preparation of this work the author(s) used ChatGPT 4.1 in order to improve language and readability. After using this tool/service, the authors reviewed and edited the content as needed and take full responsibility for the content of the publication.

## Competing Interests

AYM and JV are co-founders of Mutagentech.

## Acknowledgements

This research was supported by NIH grant 19AG056278 to JV

## Author contributions

LJN, AYM and JV conceived the project. GFH, AYM, JV, and LJN wrote the manuscript. CB and SMM collected the samples and helped with results intepretation. GFH, ML, and ELS performed the experiments. AYM and SS analyzed the data.

---

**Supplementary Figure 1.**
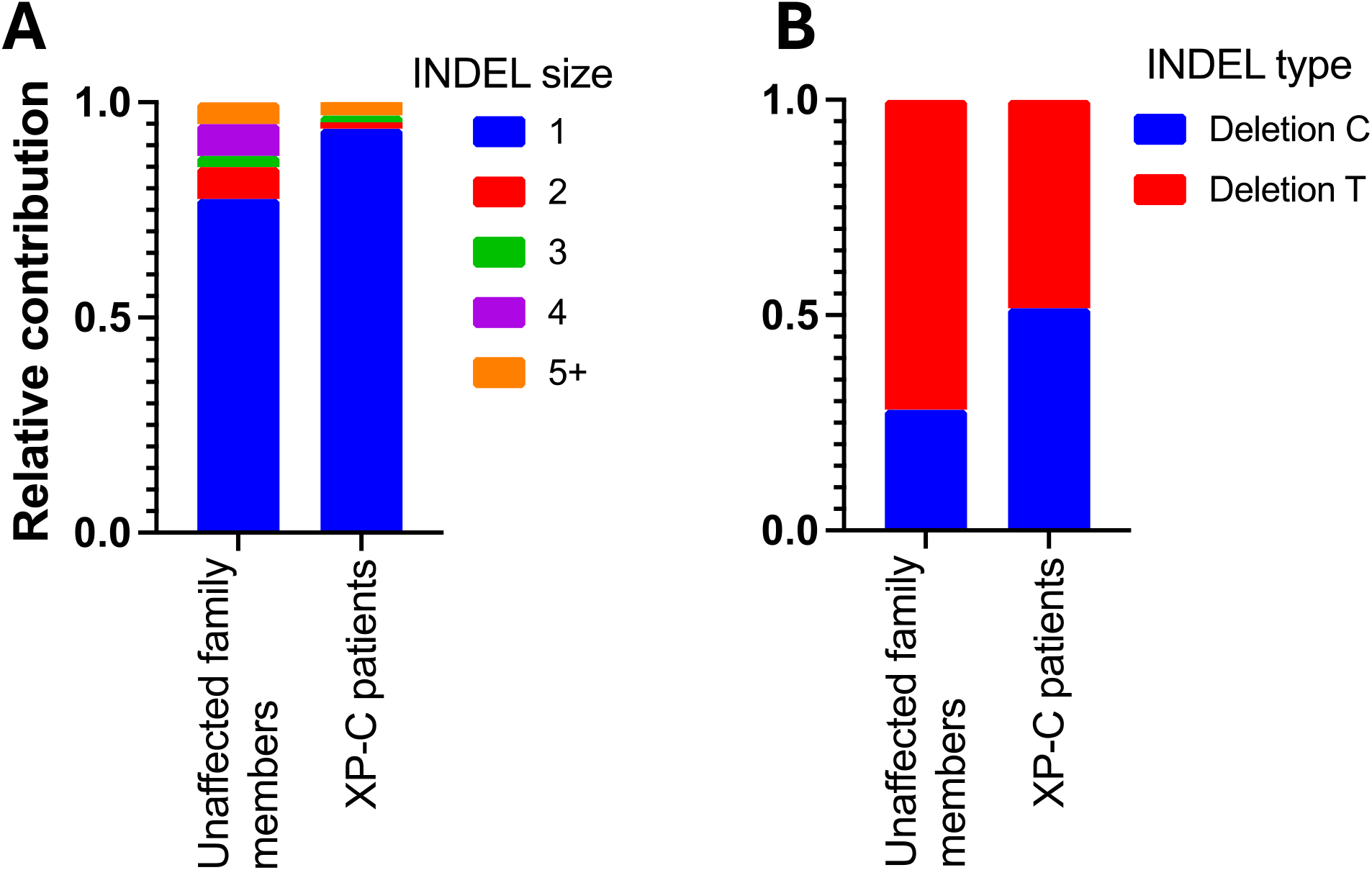
Distribution of INDELs identified in peripheral blood from XP-C patients and heterozygous carriers by size (A) and by nucleotide type for single-nucleotide deletions (B).

**Supplementary Figure 2.**
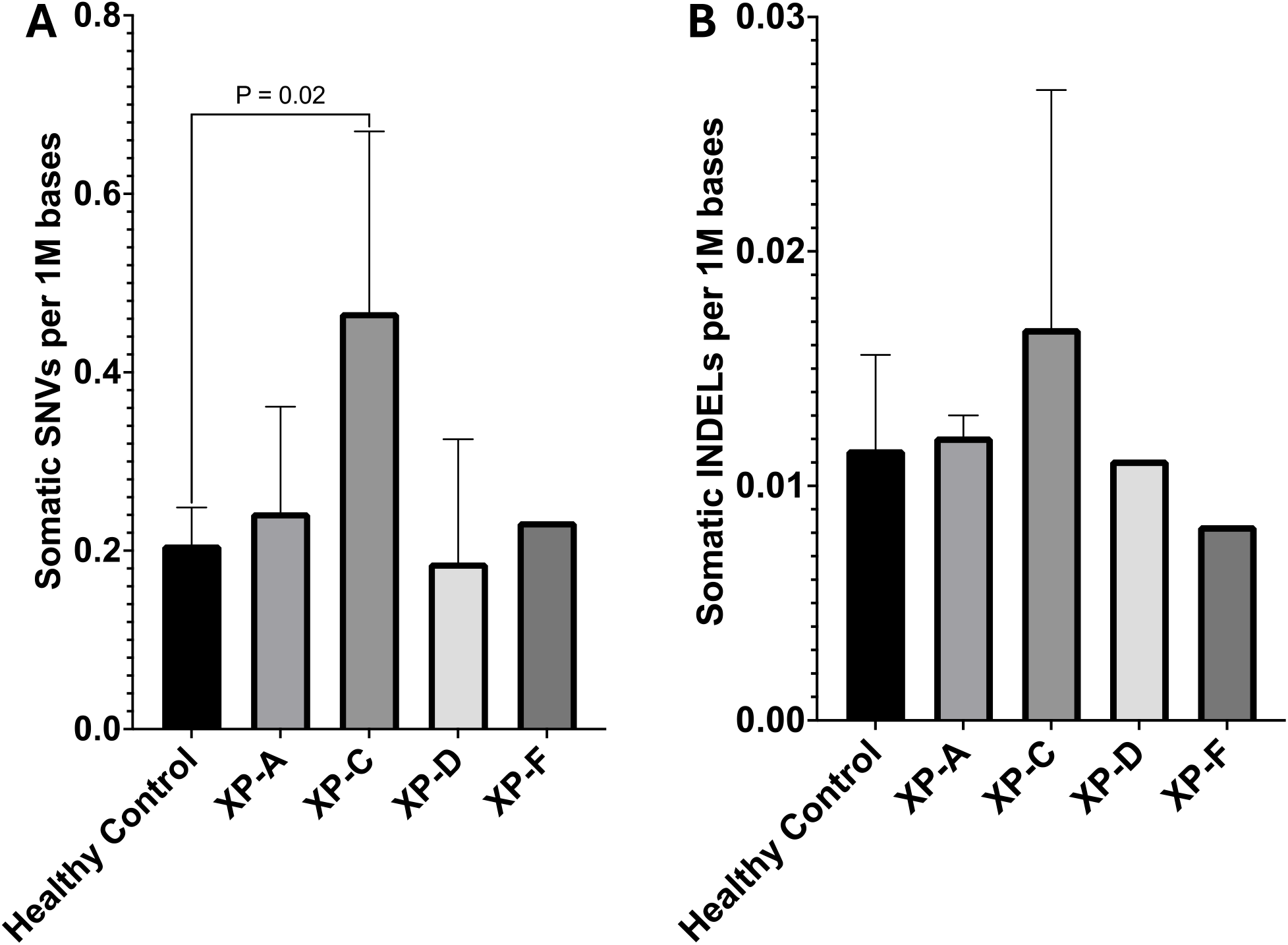
Spontaneous somatic mutation frequencies of SNVs (A) and INDELs (B) in cultured primary fibroblasts from XP patients across different complementation groups compared to fibroblasts from healthy individuals without XP-related mutations. Data are presented as mean ± SD.

**Supplementary Figure 3.**
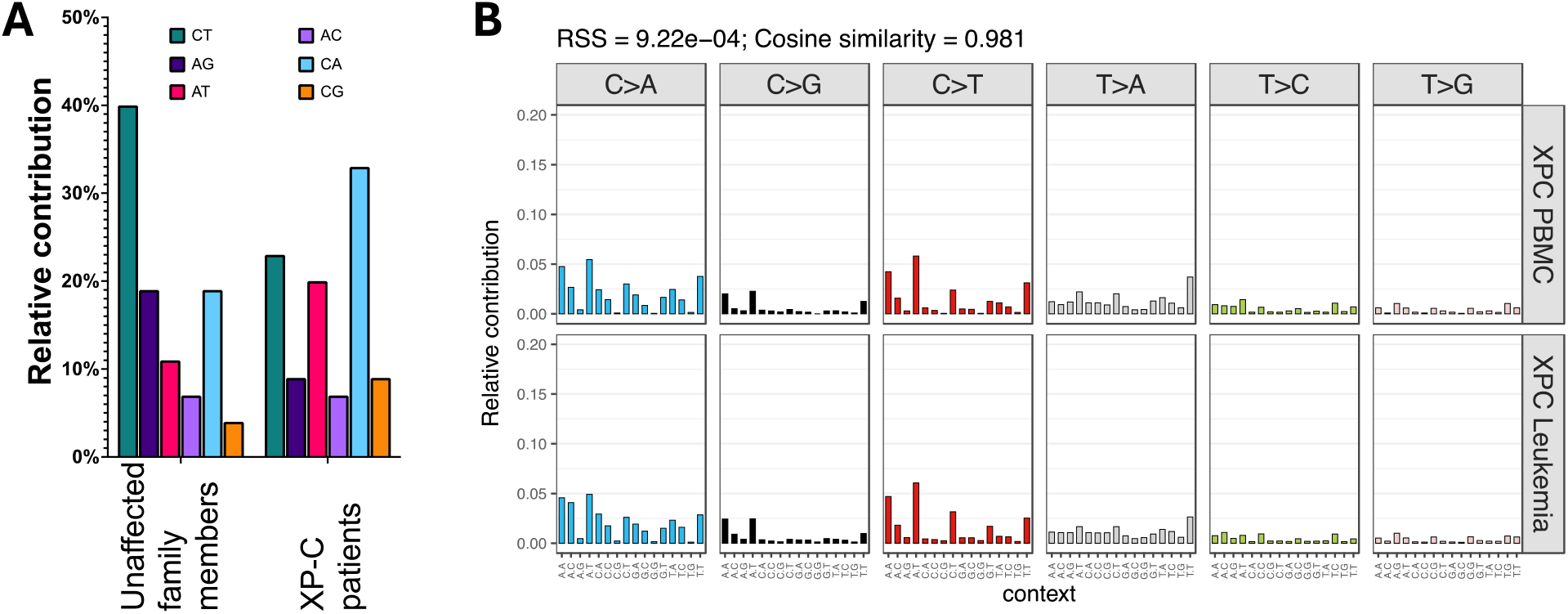
Mutation spectra in peripheral blood mononuclear cells (PBMCs) from XP-C patients. (A) Comparison of somatic mutation spectra between XP-C patients and unaffected family members, highlighting differences in substitution patterns. (B) Mutation spectra in XP-C PBMCs closely resemble those previously reported in XP-C leukemia cells (Yurchenko et al., 2020), shown as relative frequencies of each substitution type in trinucleotide contexts.

**Supplementary Table S1.** Distribution of PBMC Samples Analyzed.

| Gene | Subject | Number of subjects | Age Range |
| --- | --- | --- | --- |
| XPC | Patient | 6 | 8-24 |
|  | Parent/Sibling | 6 | 19-56 |
| XPA | Patient | 4 | 15-23 |
|  | Parent | 2 | 44-45 |
| XPD | Patient | 3 | 5-26 |
|  | Parent | 2 | 36-39 |
| XPF | Patient | 1 | 19 |
|  | Parent | 1 | 39 |

